# Dopamine depletion can be predicted by the aperiodic component of subthalamic local field potentials

**DOI:** 10.1101/2021.11.11.467452

**Authors:** Jinmo Kim, Jungmin Lee, Eunho Kim, Joon-Ho Choi, Jong-Cheol Rah, Ji-Woong Choi

**Author notes:** Corresponding authors Correspondence to: Ji-Woong Choi, Full address: E3-504, 333, Techno jungang-daero, Hyeonpung-eup, Dalseong-gun, Daegu, Republic of Korea. These authors contributed equally to this work (Jinmo Kim, Jungmin Lee).

## Abstract

Electrophysiological biomarkers reflecting the pathological activities in the basal ganglia are essential to gain an etiological understanding of Parkinson’s disease (PD) and develop a method of diagnosing and treating the disease. Previous studies that explored electrophysiological biomarkers in PD have focused mainly on oscillatory or periodic activities such as beta and gamma oscillations. Emerging evidence has suggested that the nonoscillatory, aperiodic component reflects the firing rate and synaptic current changes corresponding to cognitive and pathological states. Nevertheless, it has never been thoroughly examined whether the aperiodic component can be used as a biomarker that reflects pathological activities in the basal ganglia in PD. In this study, we examined the parameters of the aperiodic component and tested its practicality as an electrophysiological biomarker of pathological activity in PD. We found that a set of aperiodic parameters, aperiodic offset and exponent, were significantly decreased by the nigrostriatal lesion. To further prove the usefulness of the parameters as biomarkers, acute levodopa treatment reverted the aperiodic offset. We then compared the aperiodic parameters with a previously established periodic biomarker of PD, beta frequency oscillation. We found a significantly low negative correlation with beta power. We showed that the performance of the machine learning-based prediction of pathological activities in the basal ganglia can be improved by using the lowly correlated parameters, beta power and aperiodic component. We suggest that the aperiodic component will provide a more sensitive measurement to early diagnosis PD and have the potential to use as the feedback parameter for the adaptive deep brain stimulation.

## Introduction

Parkinson’s disease (PD) is the second most common neurodegenerative disorder after Alzheimer’s disease. It has been well accepted that behavioral symptoms such as bradykinesia and cognitive impairment (Collaborators, 2018; Poewe et al., 2017) are mainly attributable to some extent of dopaminergic cell death. PD brains show clear pathological hallmarks, including the loss of dopaminergic neurons in the substantia nigra pars compacta (SNc) (Poewe et al., 2017). The reported abnormal neural activity patterns in the basal ganglia (Petersson et al., 2020) and the behavioral symptoms are ameliorated by the dopaminergic medication such as dopamine precursor levodopa (L-DOPA) (Charvin et al., 2018). However, the underlying mechanisms of abnormal brain activity remain unclear. The lack of understanding makes it difficult to determine which electrophysiological features most closely reflect the pathological condition.

Since the finding that the subthalamic nucleus (STN) shows exaggerated and synchronized oscillatory activity in the beta frequency range in PD patients (Levy et al., 2000; Marsden et al., 2001), the majority of neurophysiological research in this field has postulated beta activity as a stereotypical biomarker across species from animals (Bergman et al., 1994; Haumesser et al., 2021; Mallet et al., 2008; Sharott et al., 2005) to PD patients (Brown et al., 2001; Gilron et al., 2021; Kuhn et al., 2009; Little and Brown, 2014). Abnormal beta power was shown to correlate with the clinical severity of parkinsonian motor symptoms (Neumann et al., 2016). Moreover, multiple lines of evidence have suggested that the abnormal increases in beta power are suppressed by treatment with L-DOPA (Doyle et al., 2005; Rao et al., 2006; Ray et al., 2008). However, beta frequency oscillation is far from a robust biomarker of PD. In fact, many studies in humans and animal models have reported pathological PD symptoms without significant beta oscillation (Bronte-Stewart et al., 2009; Ivica et al., 2018; Pan et al., 2016; Rosa et al., 2011). We noted the possibility that the aperiodic (nonoscillatory) component, which is the component of power spectral densities (PSD) in local field potentials (LFP) that does not include the periodic (oscillatory) component, can be a biomarker reflecting behavioral and cognitive states (Belova et al., 2021; Donoghue et al., 2020).

The aperiodic component can be described by a Lorentzian function as *L* = *b* − log(*k* + *f*^−*χ*^), where *b, k*, and *χ* represent the aperiodic offset, the knee parameter, and the aperiodic exponent (see Methods for details) (Donoghue et al., 2020). It has been hypothesized that this component reflects physiological information such as the firing rate (Manning et al., 2009), excitation-inhibition balance (Gao et al., 2017; Martin et al., 2018), and synaptic currents (Baranauskas et al., 2012; Buzsaki et al., 2012). In a clinical study, the aperiodic offset and exponent increased, assessed with magnetoencephalography (MEG) recordings in the somatosensory cortex during the resting state, in PD patients compared to controls (Vinding et al., 2021). Specifically, the slope of the broadband (exponent) decreased while patients were engaged in voluntary movements (Belova et al., 2021), and the aperiodic offset decreased in relation to cognitive states while patients were performing a visual working memory task (Donoghue et al., 2020). Although the relationship between the aperiodic component and PD exists, it has not been conclusively demonstrated that the aperiodic component can be used as a biomarker reflecting pathological activities in the basal ganglia in PD. In other words, it has not yet been demonstrated whether the aperiodic component can predict PD more accurately than the previous standard biomarker, in particular, beta power, or whether the aperiodic component can produce a synergistic effect with beta power in the prediction of PD.

To investigate whether the aperiodic component can be a useful biomarker that reflects the pathological activities in the basal ganglia, we used in vivo LFP recordings in the STN in anesthetized hemiparkinsonian rats produced by unilateral 6-hydroxydopamine (6-OHDA) lesions. Notably, we found that the parameterized aperiodic component, referred to as the aperiodic offset and exponent, were correlated with the state of dopamine (DA) depletion. The aperiodic parameters improved the performance in the machine learning-based classification of DA-depleted states when considered in conjunction with beta power. Taken together, we suggest that aperiodic parameters are useful biomarkers reflecting pathological activities in the STN in parkinsonian rodents.

## Materials and methods

### Experimental models

All experiments were approved and conducted following the guidelines of the Institutional Animal Care and Use Committee at Daegu Gyeongbuk Institute of Science and Technology (DGIST-IACUC-21051807-0001). Surgery and recording were performed in 7- to 8-week-old Sprague-Dawley male rats. The rats were housed under artificial conditions of light (12-hour light/dark cycle) with free access to food and water.

To establish unilateral 6-OHDA lesions, the rats were anesthetized using isoflurane (1.5–2.5%), followed by pretreatment with desipramine hydrochloride (25 mg/kg, i.p.; Tocris, Bristol, ENG) and pargyline (5 mg/kg, i.p.; Tocris, Bristol, ENG) to prevent noradrenergic lesions and extrasynaptic breakdown of 6-OHDA. Next, the rats were placed in the stereotaxic frame, and a craniotomy was performed. Then, 4 or 8 μl of 6-OHDA (5 μg in 1 μl saline containing 0.1% ascorbic acid; Tocris, Bristol, ENG and Sigma-Aldrich, MA, USA) was injected into the left MFB at the following coordinates: AP, −4.4 mm; ML, −1.2 mm; DV, −7.8 mm. During all surgical procedures, animals were kept warm with a handwarmer and kept on a heating pad after the surgery in an isolated cage until they regained movement.

### In vivo electrophysiology

For in vivo electrophysiological recordings, anesthesia was induced with 5% isoflurane and maintained using 1–2% isoflurane. Craniotomies were drilled to a diameter of 1 mm, leaving the dura mater intact. For grounding and referencing, two stainless steel epidural screws were placed above the ipsilateral and contralateral cerebellum. Electrophysiological recordings were performed in a Faraday cage using a neural data acquisition system (PZ5 and RZ2; Tucker-Davis Technologies, FL, USA). We dipped 16-channel NeuroNexus probes into DiI (Invitrogen, MA, USA) and slowly inserted them into the left STN (AP, −3.6 mm; ML, − 2.5 mm; DV, −7.5 mm from the dura surface), while high-pass filtered spikes were continuously monitored for fine positioning of the electrodes upon evaluation of multiunit activity as previously described.(Haumesser et al., 2017) The correct placement of the electrode was further verified by DiI tracking and microscopic evaluation. We excluded animals with inaccurately placed electrodes or trials that lacked typical STN multiunit activity.

To examine the effects of DA replacement on the expression of electrophysiological biomarkers, the rats were administered L-DOPA (7.5 mg/kg; Sigma-Aldrich, MA, USA) following baseline recordings. To assess the effects of L-DOPA, we used 1 min of data at the 30 min time points after drug administration.(Dupre et al., 2016; Giannicola et al., 2010; Ryan et al., 2018)

## Data analysis

### Calculating power spectral densities

PSD were calculated using Welch’s method (pwelch function, one 1-min recording, 2-s windows, 1-s overlap) and further analyzed using MATLAB (2020a; Mathworks, MA, USA), unless otherwise stated. Beta power was calculated using (1) the three canonical methods and (2) the FOOOF algorithm.

### Canonical methods

The following canonical methods were used to calculate beta power: (1) *broad*: the summation of PSD in the predefined broad beta band (12–35 Hz); (2) ±*2 Hz*: the summation of PSD within ±2 Hz of the maximal peak frequency in the broad beta band; and (3) *curve*: the mean PSD in the broad beta band relative to the curve fit *b* + *a* × *f*^−1.5^ on a log scale, where a and b are polynomial fitted coefficients. All summations were performed using PSD calculated by Welch’s method, which we refer to as the original PSD.

### FOOOF algorithm

We used the FOOOF algorithm which is the open-source toolbox parameterizing the original PSD into periodic and aperiodic components, developed by Voytek and colleagues (Donoghue et al., 2020). The PSD was modeled as 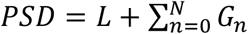, where the PSD is a combination of the aperiodic component, *L*, and *N* total Gaussians, *G*. Periodic components were parameterized as a mixture of *N* Gaussian distributions 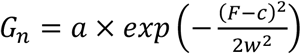, where *a* is the power of the peak on a log scale, *c* is the center frequency in Hz, and *w* is the standard deviation of the Gaussian distribution in Hz. We referred to 2*w* as the Gaussian bandwidth of *G*_*n*_ as described in the original FOOOF paper (Donoghue et al., 2020). Furthermore, the aperiodic component was also parameterized using a Lorentzian function as *L* = *b* − log(*k* + *F*^*χ*^) using the FOOOF algorithm, where *b, k*, and *χ* represent the aperiodic offset describing the y intercept, the knee parameter reflecting the bend in the aperiodic fit, and the aperiodic exponent representing the slope of the fit in log-log space when *k* = 0, respectively. The full model was described using these periodic and aperiodic parameters (fitting curves). Goodness of fit was estimated by comparing each fit with the original PSD in terms of the mean absolute error (MAE) and the *R*^2^ of the fit. For the parameterization, the 1–50-Hz frequency range was fed to the FOOOF algorithm as following reasons: (1) the usually considered upper bound of beta frequency range is certainly lower than 50-Hz (Yin et al., 2021), (2) we narrowed the upper bound of the input frequency range to avoid ambiguous frequency ranges where there may or may not be a knee, which may cause a worse fit when if we consider a knee parameter in the absence of bending in log-log space. The lower bound of the peak width limits was set as 1 Hz. For the aperiodic fit, the knee parameter *k* was ignored in this study (i.e., *k* = 0), because the averaged MAE was increased from 0.060 ± 0.018 to 0.066 ± 0.034 when we changed the aperiodic fit mode from fixed (*k* = 0) to knee (*k* ≠ 0) with restricting the maximal number of peaks as 6 preventing the overfitting of the algorithm. We evaluated the goodness of fit for all data used in this study. All other parameters were used as default values. We defined the postulated individual beta band using the maximal peak frequency *c*_*max*_ with its own Gaussian bandwidth 2*w*, where the maximal peak frequency was 12–35 Hz, which represented our predefined beta band. However, the reestablished individual beta band, i.e., *c*_*max*_ ± 2*w*, was not limited to 12 and 35 Hz. When there was no oscillatory peak within 12–35 Hz, the peak frequency, the peak power, and its Gaussian bandwidth were treated as 0.

### Statistical analysis and correlation coefficients

Statistical analyses were performed using GraphPad Prism (version 9.1.2 for Windows; GraphPad, CA, USA). Unpaired or paired *t*-tests were performed, followed by D’Agostino and Pearson normality tests. Moreover, Mann-Whitney tests were performed when one or both variables failed to pass the normality test. Since the total list of beta power (*N* = 35; control: *N* = 13; 6-OHDA: *N* = 22) did not pass the D’Agostino and Pearson normality test for the comparisons of the periodic beta power and aperiodic parameters, we calculated Spearman’s correlation coefficients using MATLAB (corr function) (Mukaka, 2012). On the other hand, we calculated Pearson’s correlation coefficients for the comparison of aperiodic parameters. All significance tests were performed as one-tailed. All data are shown as the mean ± standard error of the mean unless otherwise mentioned.

### PD Classification

To delineate the relationships of periodic beta power with the aperiodic parameters and the potential of aperiodic parameters as biomarkers reflecting pathological activities in the STN in PD, we performed classification of the DA-depleted state with the SVM using MATLAB. The SVM is a binary discriminator that detects the hyperplane that optimally separates all the data points into each class. We performed the stratified and nested cross-validation strategy to evaluate the performance of each classifier such that the total data set (control: *N* = 13; 6-OHDA: *N* = 22) was partitioned into five subsets for the outer cross-validation (Dietterich, 1998). Here, one PSD data was used for each animal so that we extracted 34 LFPs as the total data. For the 5 divided subsets, we used four subsets for training and the remaining subset for testing. Hyperparameters of each classifier were optimized using the Bayesian approach with the inner 5-fold cross-validation strategy for the four subsets (Snoek et al., 2012). This nested cross-validation was performed for each combination of feature. We performed the outer loop one hundred times to yield accurate results. Using the trained SVM models, we calculated posterior probabilities and obtained ROC curves (kfoldPredict and perfcurve function). The AUC was obtained using these ROC curves; specifically, we obtained 100 AUCs for each feature combination. All AUCs are written as mean ± standard deviation for 100 iterations.

## Data availability

The datasets and code generated during the current study are available from the corresponding author on reasonable request.

## Results

### Determination of the most precisely calculated beta power

Our purpose is to assess the usefulness of the aperiodic component separated from the PSD as the effective biomarker predicting DA depleted state in PD. We used two evaluation methods, the statistical analysis, and the machine learning technique with the support vector machine (SVM), to compare (1) the beta power calculated without separating the aperiodic component, (2) the beta power calculated with separating the aperiodic component, and (3) the aperiodic component itself (Fig. 1A). To evaluate the beta power and the aperiodic component as indicators of PD, we depleted DA by lesioning the medial forebrain bundle (MFB) with 6-OHDA and recorded the electrophysiological signals in the STN (Supplementary Fig. 1A). We found that the density of tyrosine hydroxylase-expressing (TH+) fibers was sharply reduced in the lesioned hemisphere (Supplementary Fig. 1B-C). Furthermore, significantly diminished movement, increased counterclockwise rotation, and reduced forelimb contacts were observed (Supplementary Fig. 1D-F), which validated the hemiparkinsonian rat model.

**Fig. 1.**
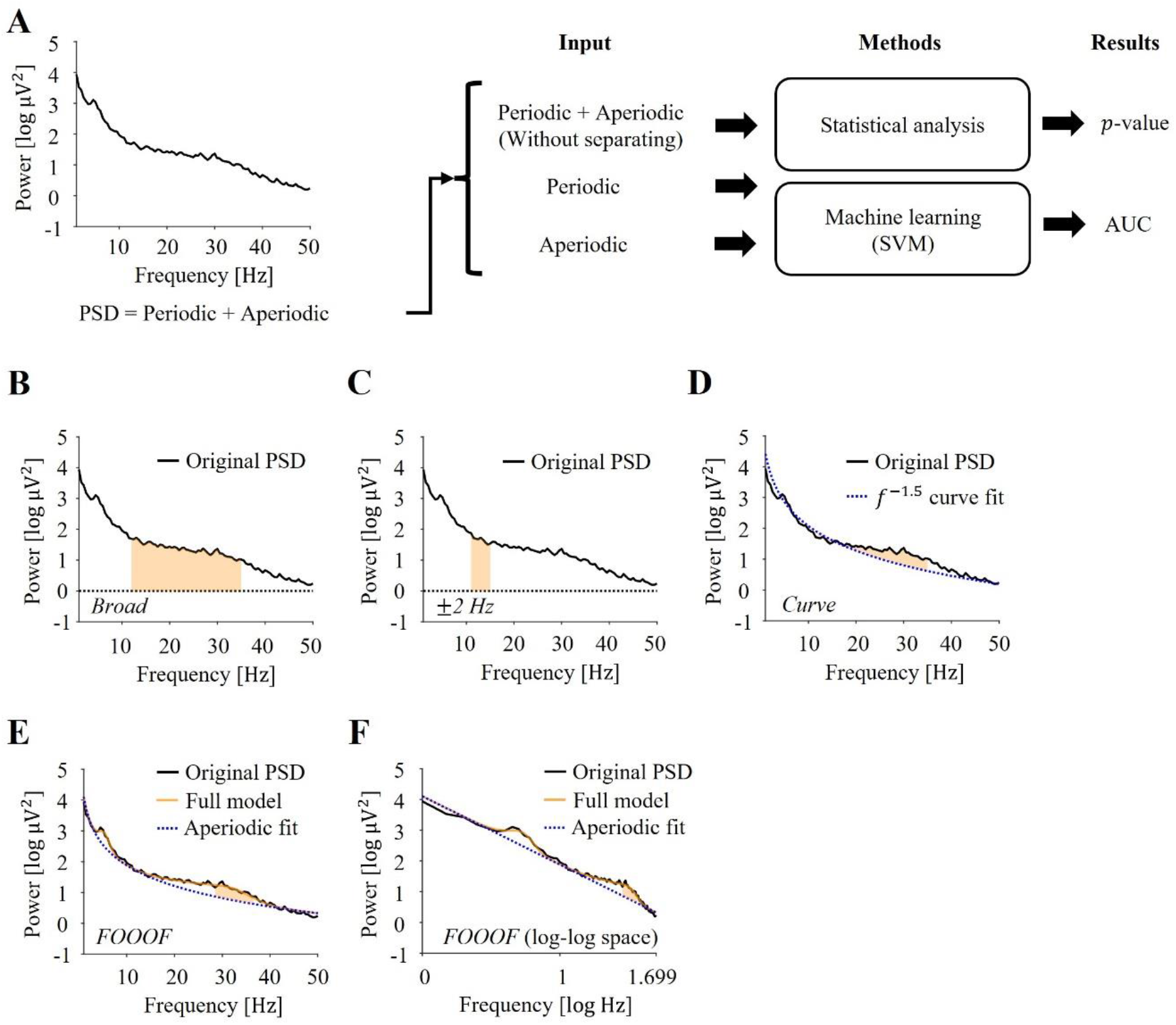
Different methods extracting electrophysiological biomarkers from the PSD. (**A**) Schematic view of what analysis we conduct in this paper. Data obtained from one 6-OHDA rat were used for this example. The original PSD denotes power spectral densities calculated using Welch’s method. Electrophysiological biomarkers extracted by different methods are evaluated using statistical analysis and machine learning with SVM. The summation of periodic and aperiodic represents that the calculated beta power without separating periodic and aperiodic components from the PSD. Two single periodic and aperiodic describe the calculated beta power with separating the aperiodic component and the aperiodic parameters obtained by the FOOOF algorithm, respectively. The performance of biomarkers is evaluated using *p*-value and the area under the receiver operating characteristic curve (AUC). (**B-F**) Examples of beta power calculation using (**B**) the *broad* method, (**C**) the ±*2 Hz* method, (**D**) the *curve* method, and (**E, F**) the FOOOF algorithm. The black solid line represents the original PSD. The shaded orange color describes spectral components used to calculate beta power. (**B**) The beta power calculation using the *broad* method (summation of the PSD in the broad beta band, 12–35 Hz). (**C**) Calculating the beta power using the ±*2 Hz* method (summation of PSD within ±2 Hz of the maximal peak frequency in the broad beta band). (**D**) The beta power calculation using *curve* method (the mean of broad beta power relative to *f*^−1.5^ curve fitted power on a log scale). The blue dotted line represents a 1/*f*^−1.5^ curve fit. (**E**) The periodic beta power calculation using the FOOOF algorithm (summation of periodic component within ±2*w* Hz around the maximal oscillatory peak), where 2*w* is the bandwidth of the Gaussian fit. The orange solid line describes the full model derived by the FOOOF algorithm. Here, the full model denotes the summation of the periodic and aperiodic fits extracted using the FOOOF algorithm. (**F**) The periodic beta power calculation using the FOOOF algorithm from the view of log-log space. The graph has the same conditions as Fig. 1E except for the scale of the *x*-axis.

First, we sought to determine the most sensitive calculation method of beta power among the currently available methods, namely, *broad*, ± *2 Hz, curve*, and the fitting oscillations and one-over-f (FOOOF) algorithm (Donoghue et al., 2020). With the exception of the FOOOF algorithm which is the open-source toolbox developed by Voytek and colleagues, the three other methods are canonical methods that calculate beta power without consideration that the aperiodic component changes according to the individual state. In this study, the original PSD denotes the PSD calculated by Welch’s method. With the *broad* and ±*2 Hz* methods, beta power was calculated by the area under the original PSD curve within the predefined beta frequency range (12–35 Hz) or the individualized beta band centered at the maximal peak with a bandwidth of ±2 Hz (Fig. 1B, C). Beta power with the *curve* method, a widely used method of calculating beta power, was obtained as follows. We fitted the *f*^−1.5^ curve to the original PSD and averaged the difference between the PSD and the *f*^−1.5^ curve across the predefined beta frequency range (Fig. 1D). The *curve* method eliminates the aperiodic component by calculating the relative PSD to the fitted curve. Since we considered that the aperiodic component can change according to PD status, we separated the periodic component from the PSD using the FOOOF algorithm, which divides spectral components into periodic and aperiodic parameters (Donoghue et al., 2020). The three separated periodic parameters, namely, the peak frequency, the peak power, and the peak bandwidth, are shown in Supplementary Fig. 2. We defined the periodic beta power as the calculated beta power within the individually postulated beta band centered at the maximal peak frequency with the peak bandwidth (Fig. 1E). Here, the full model which was the obtained fitting from the FOOOF algorithm was used instead of the original PSD to calculate the periodic beta power (see Methods for details). The term of periodic power came from that we used only the remained PSD component above the aperiodic fit which is regarded as a linear fit in log-log space (Fig. 1F).

When we defined beta power by the *broad* and ± *2 Hz* methods, no significant differences were observed in beta power between the control and 6-OHDA groups (*broad*, control: 2.695 ± 0.032 log μV^2^, *N* = 13; 6-OHDA: 2.666 ± 0.079 log μV^2^, *N* = 22; *t*_33_ = 0.276, *p* = 0.392, unpaired *t*-test; ±*2 Hz*, control: 2.531 ± 0.055 log μV^2^, *N* = 13; 6-OHDA: 2.416 ± 0.076 log μV^2^, *N* = 22; *t*_33_ = 1.060, *p* = 0.148, unpaired *t*-test; Fig. 2A, B). On the other hand, the *curve* method could successfully differentiate increased beta power by DA depletion (control: 0.047 ± 0.042 log μV^2^, *N* = 13; 6-OHDA: 0.160 ± 0.032 log μV^2^, *N* = 22; *t*_33_ = 2.165, \**p* < 0.05, unpaired *t*-test; Fig. 2C). The difference in beta power was even more significant when calculated using the FOOOF algorithm (control: 0.954 ± 0.170 log μV^2^, *N* = 13; 6-OHDA: 1.424 ± 0.073 log μV^2^, *N* = 22; *t*_33_ = 2.923, ** *p* < 0.01, unpaired *t*-test; Fig. 2D). These results suggested that the periodic beta power calculation with the FOOOF was the most sensitive method.

**Fig 2.**
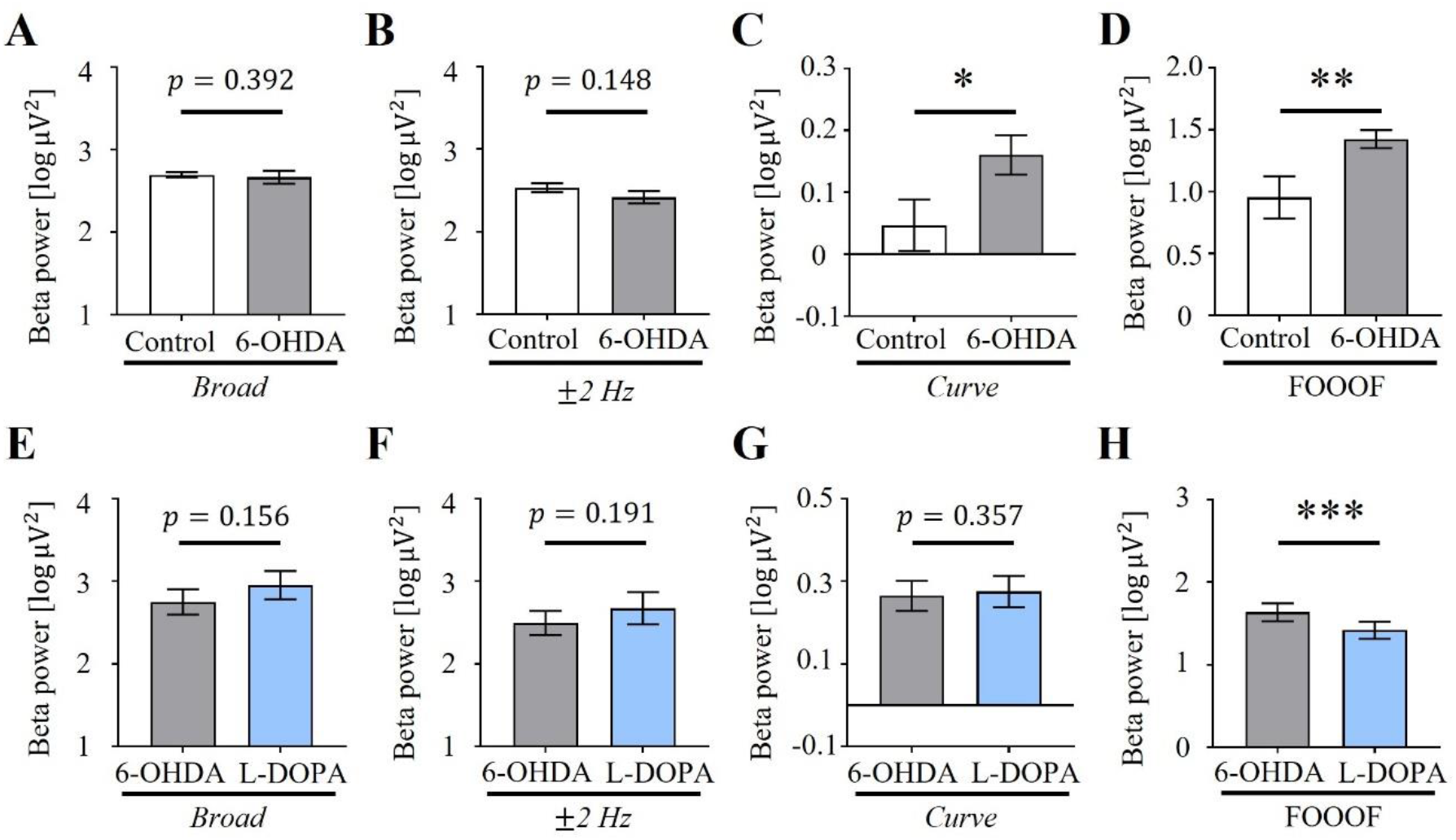
The calculated beta power with different methods. (**A-D**) Comparisons of the beta power between control and 6-OHDA groups. Error bars denote the standard error of the mean. (**A**) Canonical beta power calculation using the *broad* method (control: *N* = 13; 6-OHDA: *N* = 22; *p* = 0.392, unpaired *t*-test). (**B**) Canonical beta power calculation using the ±*2 Hz* method (control: *N* = 13; 6-OHDA: *N* = 22; *p* = 0.148, unpaired *t*-test). (**C**) Canonical beta power calculation using the *curve* method (control: *N* = 13; 6-OHDA: *N* = 22; \**p* < 0.05, unpaired *t*-test). (**D**) Periodic beta power calculation using the FOOOF algorithm (control: *N* = 13; 6-OHDA: *N* = 22; \*\**p* < 0.01, unpaired *t*-test). (**E-F**) Comparisons of the beta power before and after L-DOPA administration. Error bars denote the standard error of the mean. (**E**) Changes of the canonical beta power using the *broad* method before and after L-DOPA administration (6-OHDA: *N* = 8; L-DOPA: *N* = 8; *p* = 0.156, Wilcoxon test). (**F**) Changes of the canonical beta power using the ±*2 Hz* method before and after L-DOPA administration (6-OHDA: *N* = 8; L-DOPA: *N* = 8; *p* = 0.191, Wilcoxon test). (**G**) Changes of the canonical beta power using the *curve* method before and after L-DOPA administration (6-OHDA: *N* = 8; L-DOPA: *N* = 8; *p* = 0.357, paired *t*-test). (**H**) Changes of the periodic beta power using the FOOOF algorithm before and after L-DOPA administration (6-OHDA: *N* = 8; L-DOPA: *N* = 8; \*\*\**p* < 0.001, paired *t*-test).

To further test the sensitivity of the calculation methods, we tested whether the methods could detect a moderate yet significant DA increase following a L-DOPA injection. Because intraperitoneal (i.p.) injection of L-DOPA only partially rescued the PD-like behavioral symptoms (Supplementary Fig. 1D), the beta power difference was expected to be subtler than that between the control and 6-OHDA groups. The beta power reduced by L-DOPA was successfully detected with the periodic beta power calculation (6-OHDA: 1.635 ± 0.108 log μV^2^, *N* = 8; L-DOPA: 1.417 ± 0.104 log μV^2^, *N* = 8; *t*_7_ = 5.355, \*\*\**p* < 0.001, paired *t*-test; Fig. 2H) and not with the other three methods (*broad*, 6-OHDA: 2.752 ± 0.152 log μV^2^, *N* = 8; L-DOPA: 2.956 ± 0.170 log μV^2^, *N* = 8; *p* = 0.156, Wilcoxon test; Fig. 2E; ±*2 Hz*, 6-OHDA: 2.497 ± 0.146 log μV^2^, *N* = 8; L-DOPA: 2.676 ± 0.195 log μV^2^, *N* = 8; *p* = 0.191, Wilcoxon test; Fig. 2F; *curve*, 6-OHDA: 0.265 ± 0.037 log μV^2^, *N* = 8; L-DOPA: 0.276 ± 0.038 log μV^2^, *N* = 8; *t*_7_ = 0.381, *p* = 0.357, paired *t*-test; Fig. 2G) after the administration of L-DOPA. Furthermore, the partially rescued beta power was detected only with periodic beta power, not with the other three methods, as shown in Supplementary Fig. 3. Taken together, these results reinforced our conclusion that the periodic beta power calculation with the FOOOF determined beta power with the greatest sensitivity.

### Aperiodic parameters as novel biomarkers for PD

We then questioned whether the aperiodic parameters were reflective of pathological states in PD. We compared the aperiodic parameters in the control and 6-OHDA rats. We further examined the reversibility of the parameters by L-DOPA administration. Using the FOOOF algorithm, we obtained the aperiodic fit described as a first-order polynomial fit in log-log space (Fig. 3A, D). The aperiodic fit comprised the aperiodic offset and exponent, which represented the *y* intercept and the slope of the fit, respectively. Both the aperiodic offset and exponent were significantly lower in the 6-OHDA rats than in the control rats (offset, control: 3.792 ± 0.125 log μV^2^, *N* = 13; 6-OHDA: 3.235 ± 0.117 log μV^2^, *N* = 22; †††*p* < 0.001, Mann-Whitney test; exponent, control: 2.242 ± 0.077 μV^2^/Hz, *N* = 13; 6-OHDA: 1.968 ± 0.066 μV^2^/Hz, *N* = 22; *t*_33_ = 2.623, \*\**p* < 0.01, unpaired *t*-test; Fig. 3A-C). Furthermore, the reversibility of the decreased aperiodic offset by L-DOPA administration was also successfully detected. The aperiodic offset in 6-OHDA rats was 2.923 ± 0.218 log μV^2^ but had significantly recovered to 3.326 ± 0.171 log μV^2^ by L-DOPA (*N* = 8; *t*_7_ = 1.947, \**p* < 0.05, paired *t*-test; Fig. 3D, E). Although it was not statistically significant, the aperiodic exponent showed a tendency to increase with the administration of L-DOPA (6-OHDA: 1.771 ± 0.112 μV^2^/Hz, *N* = 8; L-DOPA: 1.883 ± 0.091 μV^2^/Hz, *N* = 8; *t*_7_ = 0.941, *p* = 0.189, paired *t*-test), as shown in Fig. 3D, F.

**Fig 3.**
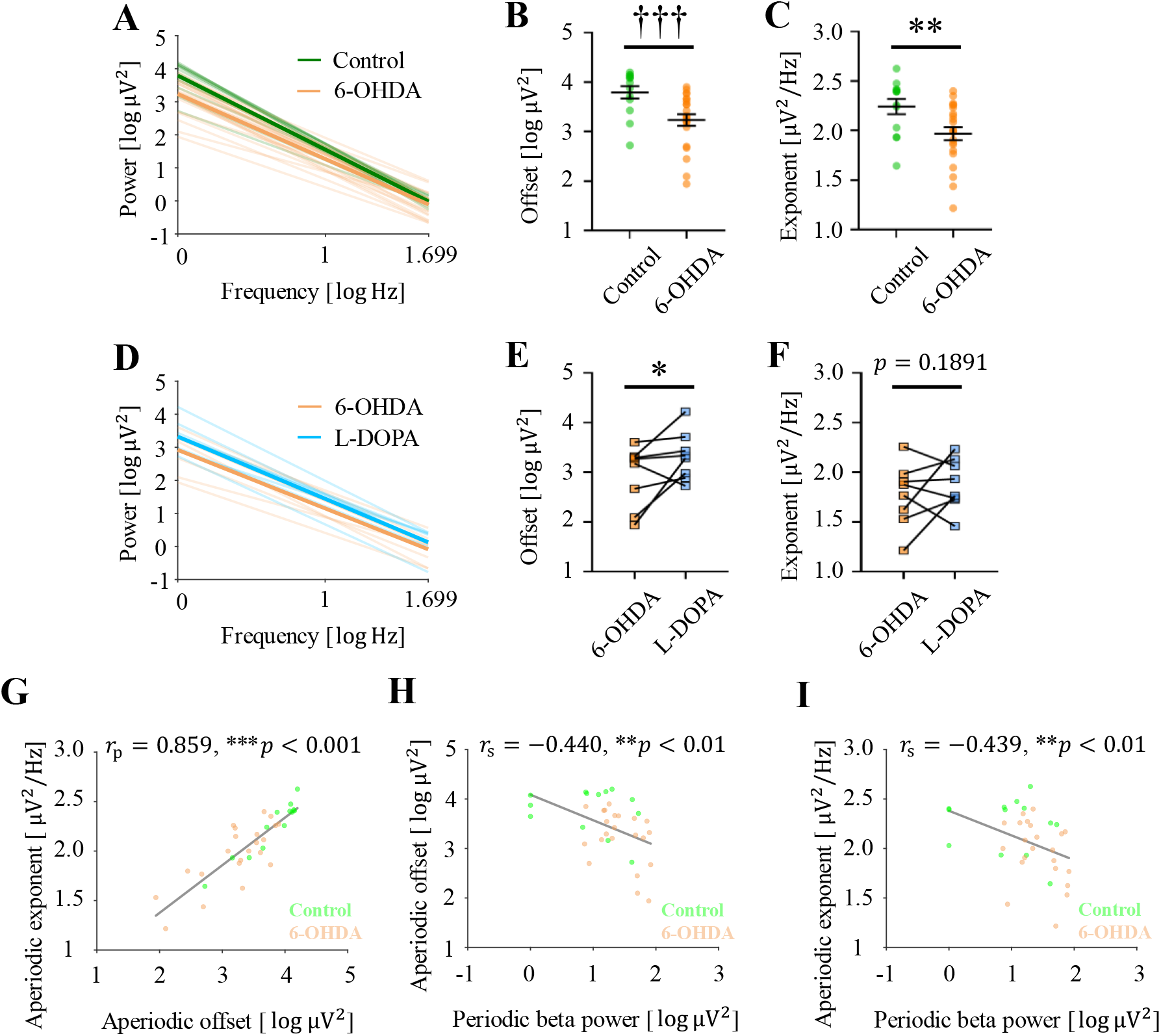
Changes of aperiodic parameters according to DA-depleted states and relationship between the periodic beta power and aperiodic parameters. (**A**) Comparison of aperiodic fit per group. For this visualization, the aperiodic fit per subject was estimated as indicated by transparent lines to reconstruct an averaged aperiodic fit for each group as shown by a thick solid line. (**B-C**) Comparisons of the aperiodic offset and exponent between control and 6-OHDA groups. Error bars denote the standard error of the mean. Green and orange dotes represent each subject. (**B**) Differences of the aperiodic offset between control and 6-OHDA groups (control: *N* = 13; 6-OHDA: *N* = 22; ††† *p* < 0.001, Mann-Whitney test). (**C**) Differences of the aperiodic exponent between control and 6-OHDA groups (control: *N* = 13; 6-OHDA: *N* = 22; \*\**p* < 0.01, unpaired *t*-test). (**D**) Comparison of aperiodic fit per group. For this visualization, the aperiodic fit per subject was estimated as indicated by transparent lines to reconstruct an averaged aperiodic fit for each group as shown by a thick solid line. (**E-F**) Comparisons of the aperiodic offset and exponent before and after L-DOPA administration. (**E**) Changes of the aperiodic offset before and after L-DOPA administration (6-OHDA: *N* = 8; L-DOPA: *N* = 8; \**p* < 0.046, paired t-test). (**F**) Changes of the aperiodic exponent before and after L-DOPA administration (6-OHDA: *N* = 8; L-DOPA: *N* = 8; *p* = 0.189, paired *t*-test). (**G**) Distributions of the periodic beta power and aperiodic parameters calculated using the FOOOF algorithm (offset-exponent: *r*_*p*_ = 0.859, \*\*\**p* < 0.001; beta-offset: *r*_*s*_ = −0.440, ** *p* < 0.01; beta-exponent: *r*_*s*_ = −0.439, ** *p* < 0.01; *r*_*p*_ and *r*_*s*_ represent the correlation coefficients of Pearson and Spearman, respectively). Green and orange colors represent the control rats (*N* = 13) and 6-OHDA rats (*N* = 22), respectively. Gray lines represent the first-order polynomial fitted lines for all data. Three control rats had zero-valued beta power.

### Aperiodic parameters provide additional information about pathological states

Having established that the aperiodic parameters change in a manner that corresponded with the depletion and replenishment of DA in vivo, we examined the possible correlation between the aperiodic parameters and traditional beta power. First, the aperiodic parameters had a significant and high positive correlation between each other (offset-exponent: *r*_*p*_ = 0.859, \*\*\**p* < 0.001, Pearson’s correlation coefficient; Fig. 3G). However, periodic beta power had low yet significant negative correlations with the aperiodic offset and with the exponent (beta-offset: *r*_*s*_ = −0.440, ** *p* < 0.01; beta-exponent: *r*_*s*_ = −0.439, ** *p* < 0.01, Spearman’s correlation coefficient; Fig. 3H, I). These results suggested that both aperiodic parameters from the FOOOF algorithm could provide additional measures that reflect pathological states.

To directly test this hypothesis, we tested whether aperiodic parameters could improve the prediction of the PD state. We performed the outer stratified 5-fold cross-validation which divides the data set (control: *N* = 13, 6-OHDA: *N* = 22) into 5 subsets, and optimized hyperparameters with Bayesian approach using the inner 5-fold cross-validation (Fig. 4A). We obtained more generalized performance of the SVM model by iterating the outer cross-validation 100 times as shown in Fig. 4B. We obtained the lowest classification performance when used the beta power with three canonical methods, *broad* (AUC= 0.656 ± 0.108), ±*2 Hz* (AUC= 0.550 ± 0.110), and *curve* (AUC= 0.529 ± 0.120). For the periodic beta power, we had slightly better performance as 0.674 ± 0.092. The aperiodic offset (AUC= 0.788 ± 0.063) and exponent (AUC= 0.707 ± 0.087) had the first and second highest AUC when considered only one single feature, respectively. When we considered the combination of the offset and exponent, AUC was slightly lower than using the single offset (AUC= 0.769 ± 0.085). The combination of three parameters, the periodic beta power, offset, and exponent had also slightly lower AUC than the single offset (AUC= 0.783 ± 0.072). However, we obtained significantly better AUC using the combination of the periodic beta power and the aperiodic exponent than using either the periodic beta power or the exponent solely (both: AUC = 0.7311 ± 0.081; both vs. periodic: *N* = 100; ††† *p* < 0.001, Mann-Whitney test; both vs. exponent: *N* = 100; †*p* < 0.05, Mann-Whitney test). Furthermore, we also obtained significantly better AUC using the combination of the periodic beta power and the aperiodic offset than using each parameter solely (both: AUC= 0.816 ± 0.060; both vs. periodic: *N* = 100; †††*p* < 0.001, Mann-Whitney test; both vs. offset: *N* = 100; \*\*\**p* < 0.001, unpaired *t*-test). Here, the combination of the periodic beta power and the offset showed the best AUC over other combinations of features. Taken together, these observations suggested that aperiodic parameters provided a better and alternative set of information to predict pathological states of PD.

**Fig. 4.**
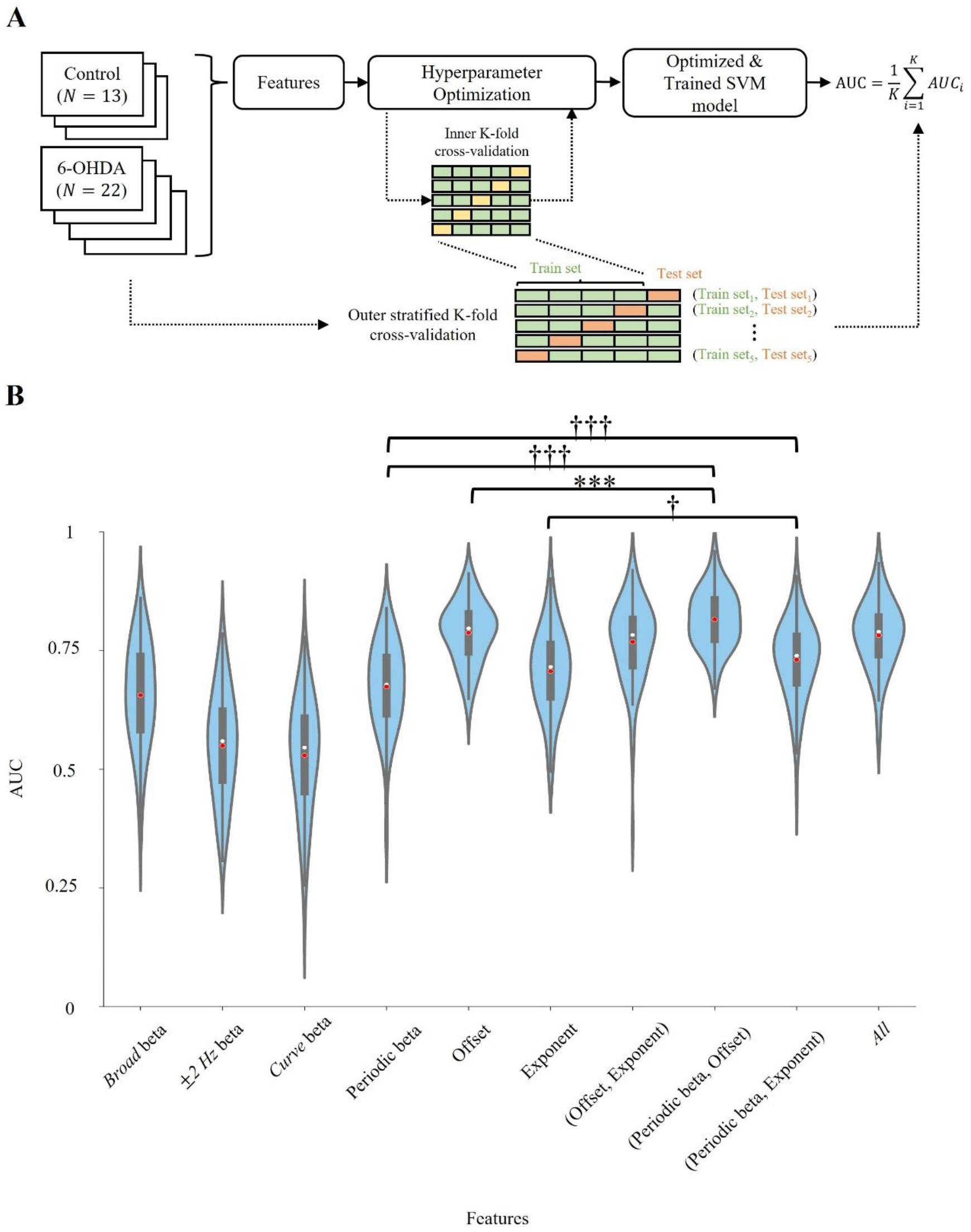
A schematic diagram for SVM model and evaluated results. (**A**) The procedure of training and evaluating SVM model. Data (control: *N* = 13; 6-OHDA: *N* = 22) with extracted features are divided into stratified *k* = 5 subsets. Train set consisted of 4 subsets and used to optimize hyperparameters with Bayesian approach. The optimization was performed with the inner *k*-fold cross-validation. The performance of optimized and trained SVM model was evaluated using the remained subset as referred to as a train set. The final performance was obtained by averaging total 5 AUCs. (**B**) Cross-validated AUCs of the SVM-based classification for PD state using different combinations of features. Periodic beta denotes the periodic beta power obtained by the FOOOF algorithm. All means the combination of the periodic beta, offset, and exponent. Violin plot shows full distributions of total 100 AUCs, where the white and red dots represent median and mean values, respectively (AUCs: *N* = 100; †*p* < 0.05, †††*p* < 0.001, Mann-Whitney test; \*\*\**p* < 0.001, unpaired *t*-test;). Moreover, thick lines show the first and third quartiles while thin lines display 95% confidence intervals.

## Discussion

Beta power is the most commonly used electrophysiological biomarker to investigate pathological activities in those with PD. Beta power was recently used as a feedback signal for adaptive deep brain stimulation (DBS) to reduce power consumption and side effects by delivering stimulation only when necessary in PD (Gilron et al., 2021; Pina-Fuentes et al., 2017). However, beta power is not robust enough to reliably predict the pathological condition associated with PD. This is known because beta power has not been exhibited in all clinical cases (Bronte-Stewart et al., 2009; Rosa et al., 2011), and there was the reemergence of tremor in some patients with the beta power-based adaptive DBS (Pina-Fuentes et al., 2020). In other words, it is necessary to extract additional information from electrophysiological recordings that reflect the state-specific information about an individual. We noted that many previous studies regarding PD biomarkers have assessed the periodic components of PSD, such as beta, gamma, and high-frequency oscillations (Petersson et al., 2020; Yin et al., 2021), but studies have not thoroughly investigated the aperiodic component.

In the present study, we investigated whether parameterized aperiodic components may serve as alternative or additional biomarkers that reflect pathological activities in the STN in PD. Although the importance of calculating beta power with separating the aperiodic component has been highlighted (Martin et al., 2018), how the separated aperiodic components change based on the pathological activity in the basal ganglia has not been thoroughly studied. Prior studies have regarded the aperiodic component in PSD as noisy variables to be corrected for or as the baseline used to normalize oscillatory power, rather than to extract them as biomarkers that reflect pathological activity in the basal ganglia. Nevertheless, several supporting pieces of evidence have suggested that the aperiodic component reflects physiological information, such as the excitation-inhibition balance, overall firing rate, and the sum of the pre- and postsynaptic currents (Baranauskas et al., 2012; Gao et al., 2017; Manning et al., 2009). A recent study also suggested that the parameterized aperiodic component is a physiological biomarker reflecting age and cognitive status (Donoghue et al., 2020). However, it has not been investigated whether these aperiodic parameters showed changes with the occurrence and alleviation of pathological activities associated with PD. In the current study, we demonstrated that the aperiodic parameters changed with the pathological states. Furthermore, the aperiodic component delivers a more robust and different set of information improving the beta power-based prediction of PD state. Taken together, we propose that aperiodic parameters could be used as practical biomarkers in PD.

### Physiological relevance of the aperiodic changes in PD animals

First, we demonstrated decreases in both the aperiodic offset and exponent in the PD animal models compared with the control animals. We believe that these decreases in aperiodic parameters showed specific pathological changes, such as altered synaptic transmission or activity patterns of the STN neurons (Buzsaki et al., 2012; Manning et al., 2009) for the following reasons.

The observed changes in the aperiodic parameters cannot be a result of different ages or task-related cognitive states (Donoghue et al., 2020; Martin et al., 2018) because animals with similar ages (7–8 weeks) were used under the same anesthetic conditions. Moreover, the decreased aperiodic offset was significantly rescued by acute L-DOPA administration. The aperiodic exponent in over half of the PD animals was also recovered by L-DOPA. These shifts in aperiodic parameters may reflect the therapeutic effect of L-DOPA, which recruits inhibitory inputs from the globus pallidus externa (GPe) to the STN, rebalancing the excitation-inhibition ratio (Belova et al., 2021). Furthermore, we found significant negative correlations between the aperiodic parameters and periodic beta power. These negative correlations implicate the traditional model of the basal ganglia network, whose elevated synchronized and phase-locked populations in PD patients lead to the minimization of neural information representation (Bar-Gad et al., 2003; Bergman et al., 1998). That is, it might be inferred that a decrease in the aperiodic component relates to more neural synchronized population in the PD state (Hammond et al., 2007; Miller et al., 2014).

### Validation of aperiodic parameters as practical biomarkers

To investigate whether aperiodic parameters can be used as practical biomarkers reflecting pathological activity in PD, we compared aperiodic parameters with the traditional electrophysiological biomarker, beta power. Since analyzing oscillatory power without separating the aperiodic component from the PSD can be misinterpreted or overstated because of a masking effect, as mentioned in the previous study (Martin et al., 2018), we calculated the periodic beta power by separating the aperiodic component from the PSD. Indeed, in our data, beta power, when including the aperiodic component, was not sensitive to the pathological changes between the control and hemi-parkinsonian animals or the reversal by L-DOPA (Fig. 2).

We found the low magnitude of the correlations between the aperiodic parameters and periodic beta power, which may imply that the aperiodic parameters and periodic beta power represent distinct sets of information about physiological or pathological activities. In other words, aperiodic parameters and periodic beta power may provide complementary information regarding an understanding of PD. To validate this idea and to evaluate whether aperiodic parameters can be used as practical biomarkers reflecting pathological activities in the STN in PD, we performed binary classification identifying the control and PD rats using the SVM classifier. We obtained a higher AUC when aperiodic parameters were adopted than when only periodic beta power was used. The lower performance when the aperiodic offset and exponent were simultaneously adopted is thought that the nonredundant (unique) information did not increase in comparison with the increased dimension since two aperiodic parameters had a high positive correlation. This is the reason that some methods of correlation-based feature selection were suggested before (Hall, 1999). On the other hand, combinations of the parameters which had the low magnitude of the correlations such as the periodic beta power and aperiodic offset or exponent had higher AUC than each parameter was used as a single feature. Taken together, our results add support to the idea that the aperiodic parameters reflect pathological activity in the STN and are potentially suitable as biomarkers of PD.

In this study, there were some limitations. Contrary to our expectation that the aperiodic offset and exponent would be increased by L-DOPA administration, only the aperiodic offset was significantly increased. The aperiodic exponent tended to increase after 6-OHDA administration in over half of the rats, but the increase was not significant. Although we performed recordings under the same anesthetic conditions in the control and PD groups to abate the impact of global suppression induced by isoflurane anesthesia, the possibility that global suppression can alter the aperiodic exponent still exists (Gao et al., 2017). Alternatively, these results may have been affected by the different origins of these two aperiodic parameters (Donoghue et al., 2020) that may be sensitive to different network effects of acute L-DOPA administration. Furthermore, the decreased aperiodic parameters in PD animals conflict with a previous result that aperiodic parameters increased, based on MEG recordings in the somatosensory cortex during the resting state, in PD patients (Vinding et al., 2021). To circumvent these discrepancies, future studies should involve neural recordings during the awake state in multiple species, such as nonhuman primates and humans. In addition, DBS effects on the aperiodic parameters are noteworthy. We expect that adaptive DBS that considers aperiodic parameters may improve DBS performance by using the complementary information about pathological activities in PD. To confirm the utility of aperiodic parameters as biomarkers for adaptive DBS (Krauss et al., 2021), we should investigate the effect of DBS on the aperiodic component in PD. Furthermore, we should overcome the computational cost caused by using the additional algorithm such as FOOOF.

The aperiodic component is a useful biomarker that can be used to better predict the pathological state of PD, rather than be considered a noisy variable to be corrected for or a baseline against which the periodic power can be normalized. We show that the aperiodic component contains a distinct set of information that reflects pathological activities and differs from the periodic component. We expect that these results will contribute to increasing the quality of diagnosing and treating the disease while furthering the etiological understanding of PD. The utility that the aperiodic component provides regarding pathological information may be worth widely investigating across neurologic and psychiatric disorders.

## Abbreviations

6-OHDA: 6-hydroxydopamine
AUC: Area under the receiver operating characteristic curve
DA: Dopamine
DBS: Deep brain stimulation
FOOOF: Fitting oscillations and one-over-f
GPe: globus pallidus externa
i.p.: intraperitoneal
L-DOPA: levodopa
LFP: Local field potentials
MEG: Magnetoencephalography
MFB: Medial forebrain bundle
PD: Parkinson’s disease
PSD: Power spectral densities
SNc: Substantia nigra pars compacta
STN: Subthalamic nucleus
SVM: Support vector machine
TH+: Tyrosine hydroxylase-expressing

## Author contributions

J.K. and J.L. equally contributed to this research. J.K., J.L., J.-C.R., and J.-W.C. designed research; J.K. developed computer code; J.K., J.L., and E.K. analyzed data; J.L. and J.K. carried out the experiment; J.L. performed the behavioral tests and immunohistochemistry staining; J.K. and J.L. wrote manuscripts in consultation with J.C., J.-C.R., and J.-W.C.

## Supplementary Methods

### Behavioral task

Behavioral tests were performed after 10 days of recovery from the first surgery. For the cylinder asymmetry test, individual rats were placed in a clean acrylic cylinder without habituation. Over 20 counts of touching the wall with fully extended digits were obtained, and the percent contralateral paw use was calculated as a ratio of total paw use. An open-field test was performed to evaluate general locomotor activities. The rats were individually placed in the center of the square chamber. Locomotor activity was recorded during the 10-min recording period through the RV2 (Tucker-Davis Technologies, FL, USA) processer and further analyzed by Ethovision 13 (Nodulus). The open-field chambers were washed with 75% ethanol solution before each behavioral testing trial to eliminate odors left by the previous rat.

### Histology

Following the recording, animals were anesthetized with 2,2,2-tribromoethanol (Sigma-Aldrich, MA, USA) and intracardially perfused with warm 1X PBS at room temperature, followed by cold 4% paraformaldehyde.

The brains were rapidly removed from the skull following decapitation and immersed in 4% paraformaldehyde for 24 h at 4°C. Subsequently, the tissues were cryoprotected with 30% sucrose. Coronal brain sections of the striatum and STN (80-μm thickness) were obtained using a cryostat. Immunostaining was performed on free-floating sections. Briefly, sections were blocked in 5% normal donkey serum in PBS (with 0.3% Triton X-100 for permeabilization) for 1-h at room temperature followed by incubation with 1:200 rabbit anti-TH antibody (AB152; Merck Millipore, MA, USA) overnight at 4°C. Staining was performed using a Vectastain Universal Elite ABC kit (Vector Laboratories, Inc. USA). Biotinylated anti-rat IgG secondary antibodies (1:200) were used to recognize primary antibodies at 37°C for 60 min, followed by washing three times and incubation with a streptavidin-horseradish peroxidase complex (1:1,000) at 37°C for 30 min. The immunoreactivities were visualized by 3,3-diaminobenzidine (DAB) within 2 min.

## Supplementary Figures

**Supplementary Fig. 1.**
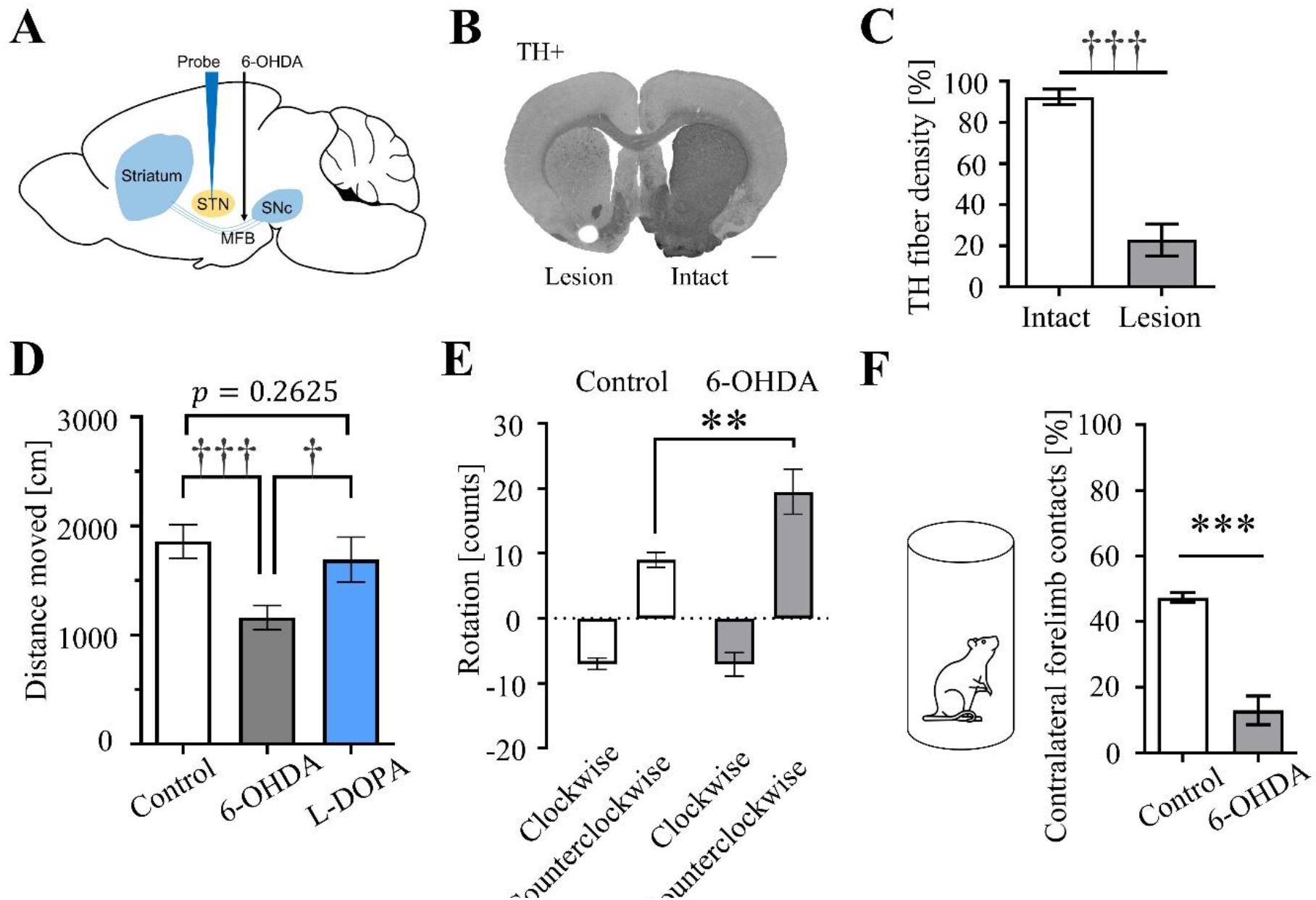
Establishment of hemiparkinsonian animal model. (**A**) Experimental scheme of the electrophysiological recording and nigrostriatal lesion. (**B**) Immunohistochemistry staining for TH+ fibers in the striatum. Scale bar=1 mm. (**C**) TH+ fiber density in the intact hemisphere and lesioned hemisphere (*N* = 7; †††*p* < 0.001, Mann-Whitney test for differences between intact and lesioned hemispheres). (**D-F**) Estimated movement of the animals in the open-field test (D, E) and cylinder test (F). (**D**) Total distance of movement in control and hemiparkinsoninan animals (control: *N* = 20; 6-OHDA: *N* = 23; L-DOPA: *N* = 4; †*p* < 0.05, †††*p* < 0.001, Mann-Whitney test for differences between control and 6-OHDA groups). (**E**) Spontaneous clockwise and counterclockwise rotations of the control and hemiparkinsoninan animal models (control: *N* = 10; 6-OHDA: *N* = 12; ** *p* < 0.01, unpaired *t*-test for differences between control and 6-OHDA groups). (**F**) Hemiparkinsonian rat in the cylinder test (left) and the percentage of contralateral forelimb contacts of the control and hemiparkinsonian rats (right; control: *N* = 13; 6-OHDA: *N* = 11; \*\*\**p* < 0.001, unpaired *t*-test for differences between control and 6-OHDA groups).

**Supplementary Fig. 2.**
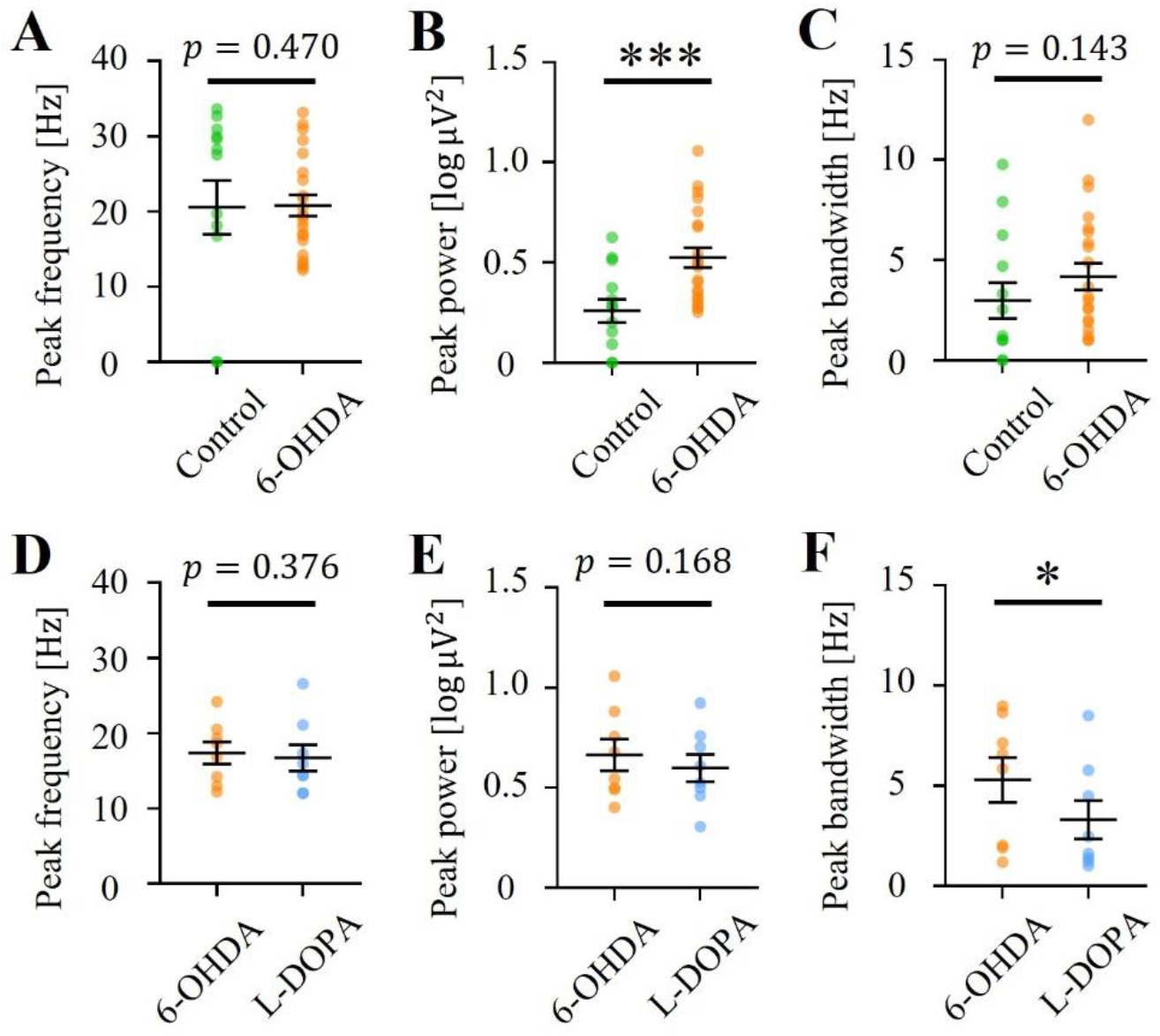
Three periodic parameters extracted by the FOOOF algorithm. (**A-C**) Comparisons of the periodic parameters between control and 6-OHDA groups. Green and orange dots represent each subject. (**A**) The extracted maximal peak frequency within the predefined beta band (12–35 Hz; control: *N* = 13; 6-OHDA: *N* = 22; *p* = 0.470, unpaired *t*-test for differences between control and 6-OHDA groups). (**B**) The maximal oscillatory peak power within the predefined beta band (control: *N* = 13; 6-OHDA: *N* = 22; \*\*\**p* < 0.001, Mann-Whitney test for differences between control and 6-OHDA groups). (**C**) The bandwidth of the maximal oscillatory peak (control: *N* = 13; 6-OHDA: *N* = 22; *p* = 0.143, unpaired *t*-test for differences between control and 6-OHDA groups). (**D-F**) Comparisons of the periodic parameters between 6-OHDA and L-DOPA groups. Orange and blue dots represent each subject. (**D**) The maximal peak frequency within the predefined beta band (6-OHDA: *N* = 8; L-DOPA: *N* = 8; *p* = 0.376, paired *t*-test for differences between 6-OHDA and L-DOPA groups). (**E**) The maximal oscillatory peak power within the predefined beta band (6-OHDA: *N* = 8; L-DOPA: *N* = 8; *p* = 0.168, paired *t*-test for differences between 6-OHDA and L-DOPA groups). (**F**) The bandwidth of the maximal oscillatory peak (6-OHDA: *N* = 8; L-DOPA: *N* = 8; * *p* < 0.05, paired *t*-test for differences between 6-OHDA and L-DOPA groups).

**Supplementary Fig. 3.**
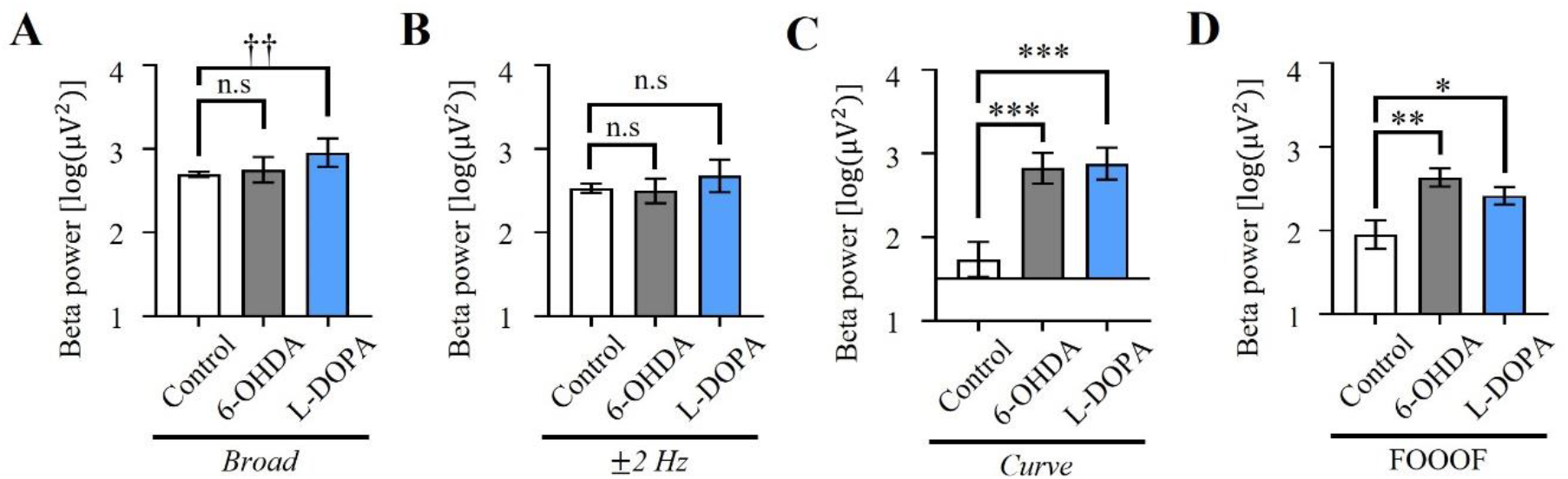
Beta power according to the DA-depleted state. (**A-D**) Comparisons of the calculated beta power between control, 6-OHDA, and L-DOPA groups for each method. (**A**) Comparisons of the beta power according to the DA-depleted state using the *broad* method (control: *N* = 13; 6-OHDA: *N* = 8; L-DOPA: *N* = 8; *p* = 0.329, unpaired *t*-test for differences between control and 6-DA groups; †† *p* < 0.01, Mann-Whitney test for significant differences between control and L-DOPA groups). (**B**) Comparisons of the beta power according to the DA-depleted state using the ±*2 Hz* method (control: *N* = 13; 6-OHDA: *N* = 8; L-DOPA: *N* = 8; *p* = 0.401, unpaired *t*-test for differences between control and 6-OHDA groups; *p* = 0.081, Mann-Whitney test for differences between control and L-DOPA groups). (**C**) Comparisons of the beta power according to the DA-depleted state using the *curve* method (control: *N* = 13; 6-OHDA: *N* = 8; L-DOPA: *N* = 8; \*\*\**p* < 0.001, unpaired *t*-test for differences between control and 6-OHDA groups; *** *p* < 0.001, unpaired *t*-test for differences between control and L-DOPA groups). (**D**) Comparisons of calculated beta power according to the DA-depleted state using the FOOOF algorithm (control: *N* = 13; 6-OHDA: *N* = 8; L-DOPA: *N* = 8; \*\**p* < 0.01, unpaired *t*-test for differences between control and 6-OHDA groups; * *p* < 0.05, unpaired *t*-test for differences between control and L-DOPA groups).

